# Gene expression analysis of the *Xenopus laevis* early limb bud proximodistal axis

**DOI:** 10.1101/2022.05.03.490399

**Authors:** D.T. Hudson, J. S. Bromell, R.C. Day, T McInnes, J.M. Ward, C.W. Beck

## Abstract

**Background:** Limb buds develop as bilateral outgrowths of the lateral plate mesoderm and are patterned along three axes. Current models of proximal to distal patterning of early amniote limb buds suggest that two signals, a distal organising signal from the apical epithelial ridge (AER, Fgfs) and an opposing proximal (retinoic acid) act early on pattern this axis.

**Results:** Transcriptional analysis of stage 51 *Xenopus laevis* hindlimb buds sectioned along the proximal-distal axis, showed that the distal region is distinct from the rest of the limb. Expression of *capn8*.*3*, a novel calpain, was located in cells immediately flanking the AER. The Wnt antagonist Dkk1 was AER-specific in *Xenopus* limbs. Two transcription factors, *sall1* and *zic5*, were expressed in distal mesenchyme. *Zic*5 has no described association with limb development. We also describe expression of two proximal genes, *gata5* and *tnn*, not previously associated with limb development. Differentially expressed genes were associated with Fgf, Wnt and retinoic acid (RA) signalling as well as differential cell adhesion and proliferation.

**Conclusions:** We identify new candidate genes for early proximodistal limb patterning. Our analysis of RA-regulated genes supports a role for transient RA gradients in early limb bud in proximal-to-distal patterning in this anamniote model organism.

## 1. Introduction

Limb bud development and patterning is best understood in amniotes (recently reviewed in McQueen and Towers, 2020). Limbs develop at specific points along the anterior-posterior axis of the embryo, and are comprised of mesenchymal cells surrounded by an epithelial layer. The mesenchymal cells are derived from lateral plate mesenchyme, and contribute the cartilage, dermis, tendons and ligaments of the limb (Pearse et al., 2007). Muscle and nerve cells migrate in from the dermamyotome and spinal cord respectively, and as such are not directly derived from the limb buds. The early limb bud has three axes which are patterned by signalling centres: proximal-distal, anterior-posterior and dorsal-ventral. The proximal to distal axis can be seen in the fore- and hindlimb limb skeletal elements of the stylopod (humerus/femur), zeugopod (radius and ulna/tibia and fibula) and autopod (wrist and hand/ankle and foot).

A key player in proximodistal patterning is a region of epithelial cells established at the boundary between dorsal and ventral limb bud surfaces (the future back and palm of the autopod). These cells establish a signalling centre, the “apical epithelial ridge” or AER, originally demonstrated by the extirpation experiments of Saunders in chickens. Saunders found that the removal of the AER at three days of development resulted in failure of the zeugopod and autopod to form, with the limb truncated at the elbow/knee level. Later AER removal resulted in a failure of just the autopod to form, suggesting that limb skeletal elements are formed in a proximal to distal order (Saunders, 1948). There are two main models for how this patterning is established. The first is the “progress zone” model (Summerbell et al., 1973), which proposes that the mesoderm closest to the AER determines cell fate. As the limb bud elongates, the first cells to leave the progress zone, or the influence of the AER, differentiate into the proximal limb (stylopod), with later cells falling out of range forming progressively more distal elements. An alternative model proposes two signals, Fgfs from the AER antagonising a proximal retinoic acid (RA) signal (Mercader et al., 2000). Supporting this “two-signal” model, the Meis homeobox transcription factors Meis1 and 2 are both RA inducible and expressed proximally in developing limb buds (Mercader et al., 2000). A more recent model, termed “signal-time” has been proposed from grafting experiments in chicken limb buds (Saiz-Lopez et al., 2015). This combines an early two signal Fgf/RA model specifying the proximal limb and transition from stylopod to zeugopod, with an intrinsic, timing dependent mechanism taking over to specify the zeugopod to autopod transition and autopod patterning.

Amphibian limb buds develop quite late compared to those of amniotes. *Xenopus laevis* tadpole hindlimbs become capable of autonomous development at stage 48, just as the tadpole begins to feed and when they are still relatively small (reviewed in Keenan and Beck, 2016). Fate maps of the hindlimb bud mesenchyme indicate that there is progressive determination of distal skeletal elements as the limb bud grows (Tschumi, 1957). Early on, at stages 48 and 49, only the pelvis and femur (stylopod) are determined, the cells destined to form the zeugopod (tibia fibula) first appear distally at stage 50 and the more proximal elements of the autopod (tarsus) at stage 51. Development of the more distal elements depends on the overlying epithelial cells. A morphological AER can be seen in sections from stage 50, but becomes indistinct by stage 53. The biological AER, *fgf8* expression in the innermost layer of the distal epidermis, persists until stage 54 where it associates with growing digits (Christen and Slack, 1997, Endo et al., 2000, Jones et al., 2013, Wang and Beck, 2014). In addition to patterning by distal *fgf8*, we have previously suggested that an opposing and transient retinoic acid gradient exists across the stage 51 *Xenopus* limb bud mesenchyme proximodistal axis. From stage 50, *aldh1a2*, the rate limiting step in RA synthesis from vitamin A, is expressed proximally, and *cyp26b*, a catabolic enzyme that degrades RA, is expressed distally in the mesenchyme from stage 51 (McEwan et al., 2011).

In this study, we set out to map differential expression by comparing transcriptomes across three equal proximodistal segments of the stage 51 *Xenopus laevis* hindlimb bud (Figure 1A). In particular, we sought evidence for differential expression of RA-regulated genes across the proximodistal axis, indicative of a transient morphogen gradient at this stage. While the limb bud at stage 51 appears externally featureless, Tschumi’s fate maps indicate that the future skeletal elements from pelvis to tarsus are already determined (Figure 1B). However, Tschumi’s specification maps show that in the absence of the epidermis, no elements of the autopod develop from grafts of the stage 51 limb bud mesenchyme (Tschumi, 1957), supporting the idea that distal elements require signals from the AER in *Xenopus*. The AER with *fgf8* expression is established (Christen and Slack, 1997) and mesenchymal retinoic acid is expected to be produced proximally and broken down distally (McEwan et al., 2011). Also in the mesenchyme, cartilage condensation of the future femur is visible histologically, although cartilage marker *sox9* is not yet expressed (Tarin and Sturdee, 1971, Miura et al., 2008, Satoh et al., 2005). The distal-most blood vessel, the marginal sinus, which sits right under the AER, is forming at this stage (Tarin and Sturdee, 1971). Nerve invasion into the limb is just beginning proximally (Tarin and Sturdee, 1971) and muscle migration has not yet commenced (Martin and Harland, 2006).

**Figure 1:**
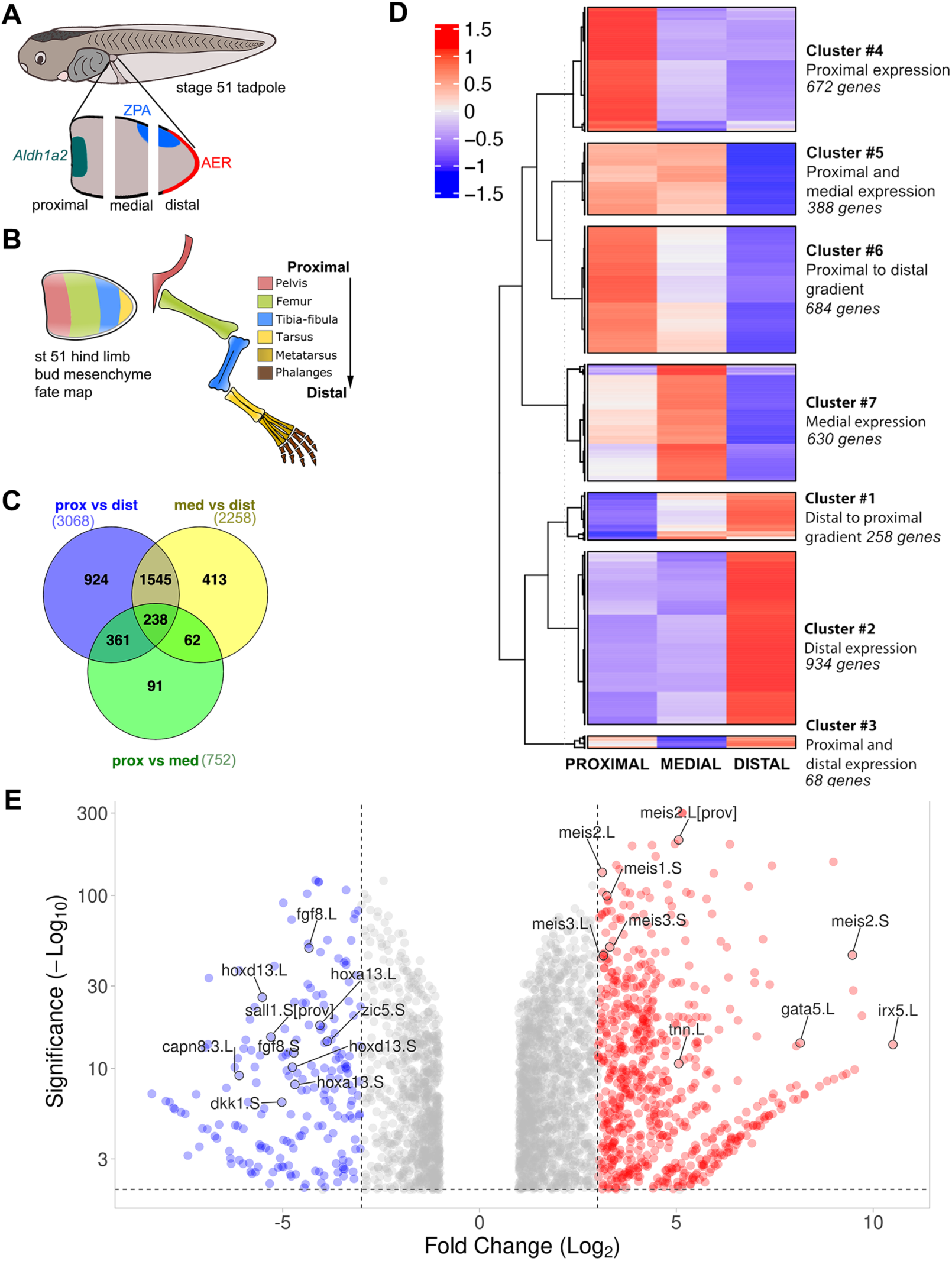
Transcriptome analysis of three regions of the early *Xenopus laevis* hindlimb bud proximal to distal axis. **A)** Schematic of sample preparation. Stage 51 hindlimb buds were divided into three equal sections along the proximal to distal axis, each part is approximately 200 μM wide. Approximate locations of the apical ectodermal ridge (AER) and zone of polarising activity (ZPA) are indicated in red and blue respectively. Location of a potential proximal source of the morphogen retinoic acid is indicated by expression of *Aldh1a2* in green. **B)** Fate map showing the contribution of developing stage 51 *X. laevis* hindlimb bud mesenchyme to skeletal elements of the limb, according to (Tschumi, 1957). **C)** Venn diagram of all significantly differentially expressed genes using Venny 2.0 (Oliveros, 2007-2015). **D)** Dendrogram heatmap showing hierarchical clustering of the differentially expressed genes, selecting for seven clusters. **E)** Volcano plot showing the relative distributions of known proximal (red: *meis1, meis2*) and distal (blue: *fgf8, hoxa13, hoxd13*) limb genes as well as the genes verified by whole mount in situ hybridisation in Figures 2,3 and 4). Genes with log2 fold change <3 in either direction are represented by grey dots.

Here, we show that genes involved with cell adhesion, cell proliferation, and three signalling pathways: Wnt, Fgf and retinoic acid, are differentially regulated in association with proximodistal position in these early *Xenopus* limb buds. Furthermore, we identify new genes associated with proximal patterning (*tnn*.*L, gata5*.*L*), and distal epithelium (*capn8*.*3*.*L*). Finally, by examining genes previously shown to be regulated by retinoic acid in salamander limb regeneration (Nguyen et al., 2017), we identify several retinoic acid-regulated genes with clearly graded expression across the proximodistal axis, supporting a role for retinoic acid in establishing this axis.

## 2. Results and Discussion

### 2.1 Transcriptome analysis of three regions of the early *Xenopus laevis* hindlimb bud proximal to distal axis identified seven clusters of differentially expressed genes

Limb buds at stage 51 were dissected into three equal parts, along the proximodistal axis, and rapidly frozen on dry ice before extracting RNA. The approximate locations of the cuts are shown in Figure 1A, with each segment measuring approximately 200 μm along the proximal-distal axis, 400 μm across the anterior-posterior axis, and 200 μm along the dorsal-ventral axis. These dissections are not expected to neatly divide the limb bud into future stylopod, zeugopod and autopod. Compared to the mesodermal fate map for this stage, the future autopod and some of the zeugopod would be derived from the distal segment, pelvis and some stylopod from the proximal segment, and stylopod/zeugopod from the medial part (Tschumi, 1957, Figure 1B). The AER is expected to be exclusively captured in distal sections, with the ZPA split between distal and medial sections. Regression analysis of mapped transcriptome read counts showed that replicate samples clustered closely together (r=0.98, 0.99 and 1.00 for proximal, medial and distal respectively). Pearson correlations of read count means showed that proximal and medial stage 51 limb bud transcriptomes are more similar to each other (r=0.97) than to distal (r=0.80 to proximal and r=0.65 to medial).

Differentially expressed genes were defined as genes with normalised read counts >log2 fold change +/-1 or higher, and with a *p* val<0.05, between any two limb bud regions (Figure 1C,D), which captured a total of 3682 genes. As expected, the number of differentially expressed genes between proximal and distal was highest (Figure 1C). Clustering of the 3682 differentially expressed genes into seven clusters best represented the patterns expected across developing limbs (Figure 1D). Cluster 2 is the largest group, containing 934 genes with predominantly distal location, including both homeologs of the AER marker *fgf8*, the ZPA morphogen *shh*.*L* and distal mesenchyme markers *hoxd13* and *hoxa13*. Cluster 4 contains 672 proximally located genes, including established proximal limb markers *meis1* and *meis2*. More graded expression is seen by genes in clusters 6 (highest proximal, 684 genes) and 1 (highest distal, 258 genes). Cluster 5 has genes that are high in proximal and medial but low distal (388 genes). Cluster 3 is the smallest, with genes that are predominantly expressed in the distal and proximal regions but not the medial (68 genes), and cluster 7 genes are opposite, with highest levels in the medial limb bud (630 genes).

### 2.2 Genes expressed in the proximal limb bud include *gata5*.*L, irx5*.*L* and *tnn*.*L*

Figure1 E shows the most proximal (red circles) genes, taking account of both fold change and significance, generated from the proximal vs. distal reads. Many of these genes are found in cluster 4 (Figure 1D), which groups 672 genes with much higher expression in the proximal limb bud than in either medial or distal sections. This group includes *aldh1a2*.*L*, formerly known as *raldh2*, previously shown to be proximally expressed by us (McEwan et al., 2011). It also includes several family members of the Meis homeobox transcription factors (*meis3*.*L, meis3*.*S, meis2*.*S, meis2*.*L, meis1*.*S*). *Meis1* and *-2* are proximal genes with a well-established functional role in limb patterning in amniotes (Delgado et al., 2020) but *meis3* has not been previously associated with limb development.

Three almost exclusively proximal genes found in Cluster 4 were verified using whole mount in situ hybridisation: *gata5*.*L, irx5*.*L* and *tnn*.*L* (Figure 2). *Gata5* encodes a zinc finger transcription factor associated with heart and liver development in *Xenopus* (Haworth et al., 2008) and has not been previously associated with limb development. *Gata5* reads were almost exclusively in the proximal limb transcriptome (Proximal>Distal Log2 FC 8.1 and 8.0 for L and S forms respectively), and whole mount *in situ* hybridisation confirmed a faint but very localised basal spot of *gata5*.*L* expression in stage 50 and 51 limb buds, which became clearer at stage 52 (Figure 2 A-C). The related transcription factor Gata6 regulates shh in anterior mouse limb development (Kozhemyakina et al., 2014,,Hayashi et al., 2016). However, this does not fit with the central basal location of *gata5*.*L* in frogs. *Gata5*.*L* in embryos (data not shown) showed clear expression in the heart as previously reported (Jiang and Evans, 1996).

**Figure 2:**
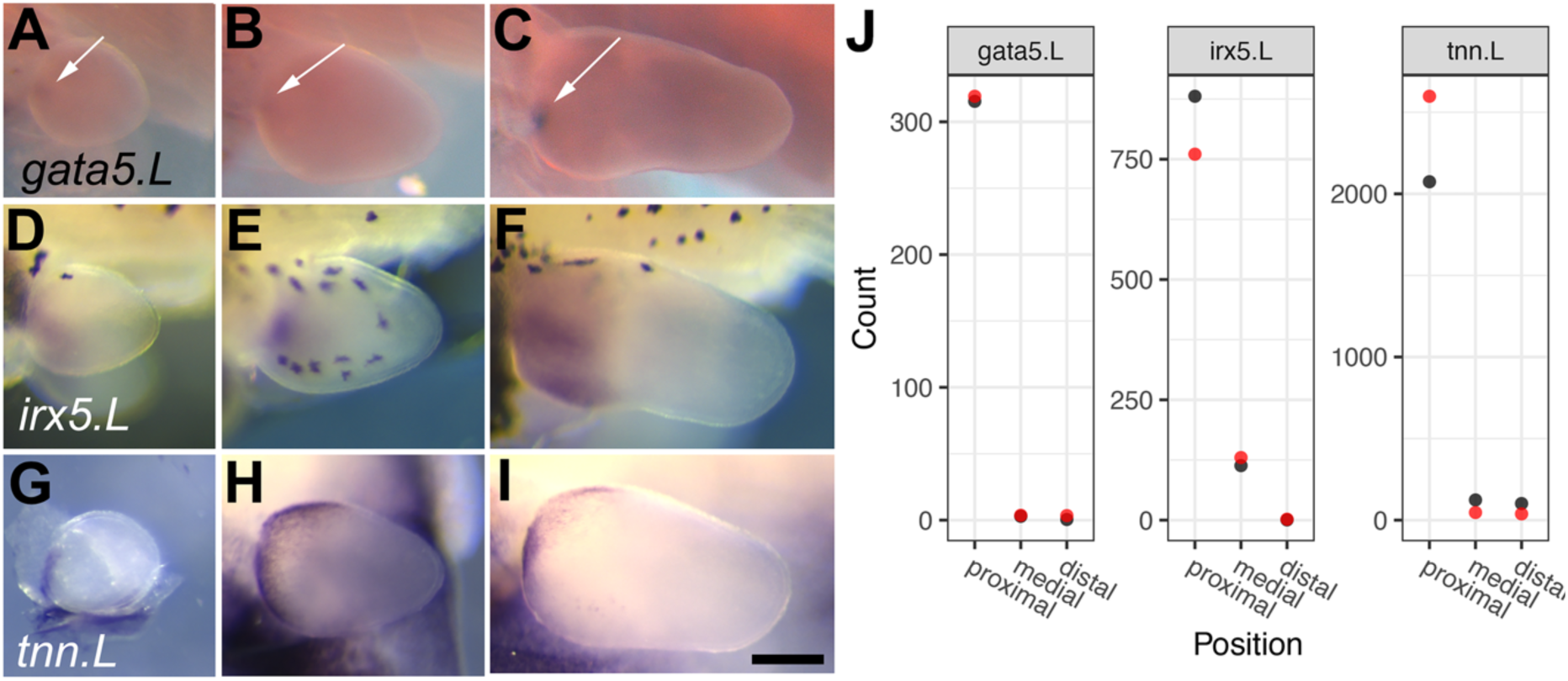
Expression of three cluster 4 proximal genes in *X. laevis* hind limb buds. **A-I** Whole mount in situ hybridisation (purple staining) of st 50-52 hind limb buds as viewed from the dorsal side, posterior uppermost. **A-C** white arrows indicate proximal spot of mesenchymal *gata5*.*L* at stage 50 (A), 51 (B) and 52 (C). **D-F** *irx5*.*L* in the proximal mesenchyme at stage 50 (D), 51 (E) and 52 (F). Black stellate cells are melanophores in non-albino tadpoles (D-F), expression in stage 52 is anterior as well as proximal. **G-I** Expression of *tnn*.*L* is very strong along the boundary with the body wall, and is epithelial. Stage 50 limb bud dissected to enable visualisation of limb bud, as there is strong expression in the ventral fin as well as the limb bud (G), stage 51 (H) and 52 (I). Scale bar in I is 250μM and applies to all panels. **J**, Normalised read counts of *gata5*.*L, irx5*.*L* and *tnn*.*L* in the three sections of the stage 51 limb bud transcriptome. Red and black dots indicate biological replicate data, where only red can be seen it overlaps completely with black.

*Irx5* encodes a homeobox transcription factor of the Iroquois family. Both L and S forms are expressed at much higher levels in proximal than in distal or medial limb segments, (P>D Log2 FC 10.5 for *irx5*.*L* and 6.8 for *irx5*.*S*). *Irx5*.*L* was the most over-represented proximal gene and in situ hybridisation for *irx5*.*L* confirmed this distribution, with expression basal, proximal and increasingly anteriorly localised in hindlimb buds from stage 50 to 52 (Figure 2 D-F). Embryonic expression of *irx5*.*L* in the brain and eye (data not shown) matched that of previous reports (Rodriguez-Seguel et al., 2009), although at this stage the *S* homeolog has the strongest expression (Session et al., 2016). *Irx5* has been previously associated with limb patterning in mammals, where it has shown it to be functionally redundant with *irx3*. Mouse double *irx3/5* knockouts have a reduced femur and are missing the tibia and digit 1 (Li et al., 2014), showing that these genes are important for both proximal and anterior limb development. *Irx3*.*L* and *S* genes show similar expression profiles to *irx5*, with P>D log2 fold change 4.8 for the L form and 6.8 for the S form, suggesting that they may also act together in *Xenopus* limb patterning. In humans, Hamamy syndrome (OMIM 611174, *IRX5*, Bonnard et al., 2012) has no reported limb defect, possibly due to compensation by *IRX3*.

Tenascins are extracellular matrix glycoproteins, and *Xenopus laevis* has a single copy of Tenascin N (*tnn*.*L*) which is expressed most strongly in the proximal third of the limb bud at stage 51, with almost no read counts in the medial or distal segments (Proximal>Distal Log2 fold change 5.1). Tenascin N was originally identified in zebrafish (Weber et al., 1998) and is more commonly known as Tenascin-W. It is a chordate-specific gene with no known links to limb development (Tucker and Degen, 2019). *Tnn*.*L* expression was confirmed as proximal in stage 50 to 52 limb buds (Figure 2 G-I) where it appeared to be epithelial, expressed where the limb bud meets the trunk. Expression persists in older limbs but the proximal localisation is lost (data not shown). In tailbud stage embryos, *tnn*.*L* was expressed only in the anatomical tail bud (data not shown).

### 2.3 Genes expressed in the distal limb bud

Distally expressed genes, with much lower levels of expression in either medial or proximal limb bud thirds are found in cluster 2, which contains 934 genes (Figure 1D). *Fgf8, bmp4, dlx2, dlx3*, and *sp8* all have localised expression in the developing AER of mouse limbs (Ahn et al., 2001, Bell et al., 2003, Bulfone et al., 1993, Gorivodsky and Lonai, 2003). Neither *bmp4*.*L nor S* were differentially expressed in *Xenopus* stage 51 hindlimbs, but *fgf8*.*S, fgf8*.*L, sp8*.*L, sp8*.*S* and the distal-less homeobox genes, *dlx2*.*S, dlx2*.*L, dlx3*.*S, dlx3*.*L*, as well as the S homeolog of *dlx4*, were all cluster 2 genes. *Bmp2*.*S* and *bmp7*.*1*.*L* also appear in this cluster.

Three genes chosen for their almost exclusive expression in the distal stage 51 limb bud were confirmed as being located exclusively in the distal epithelium (Figure 3). Their distribution was highly significantly enriched in distal stage 51 limb bud, similar to that of the AER specific gene *fgf8* (Figure 1E). *Capn8*.*3*.*L* (D>P Log2 fold change 6.2, cluster 2) encodes a calcium dependent cysteine protease (calpain). Whole mount in situ hybridisation revealed *capn8*.*3*.*L* expression in the region of the forming AER at stage 50 and 51, with expression margins becoming very defined at stage 52 and maintained in the distal epithelium of the forming digits until stage 55 at least (Figure 3 A-E).

**Figure 3:**
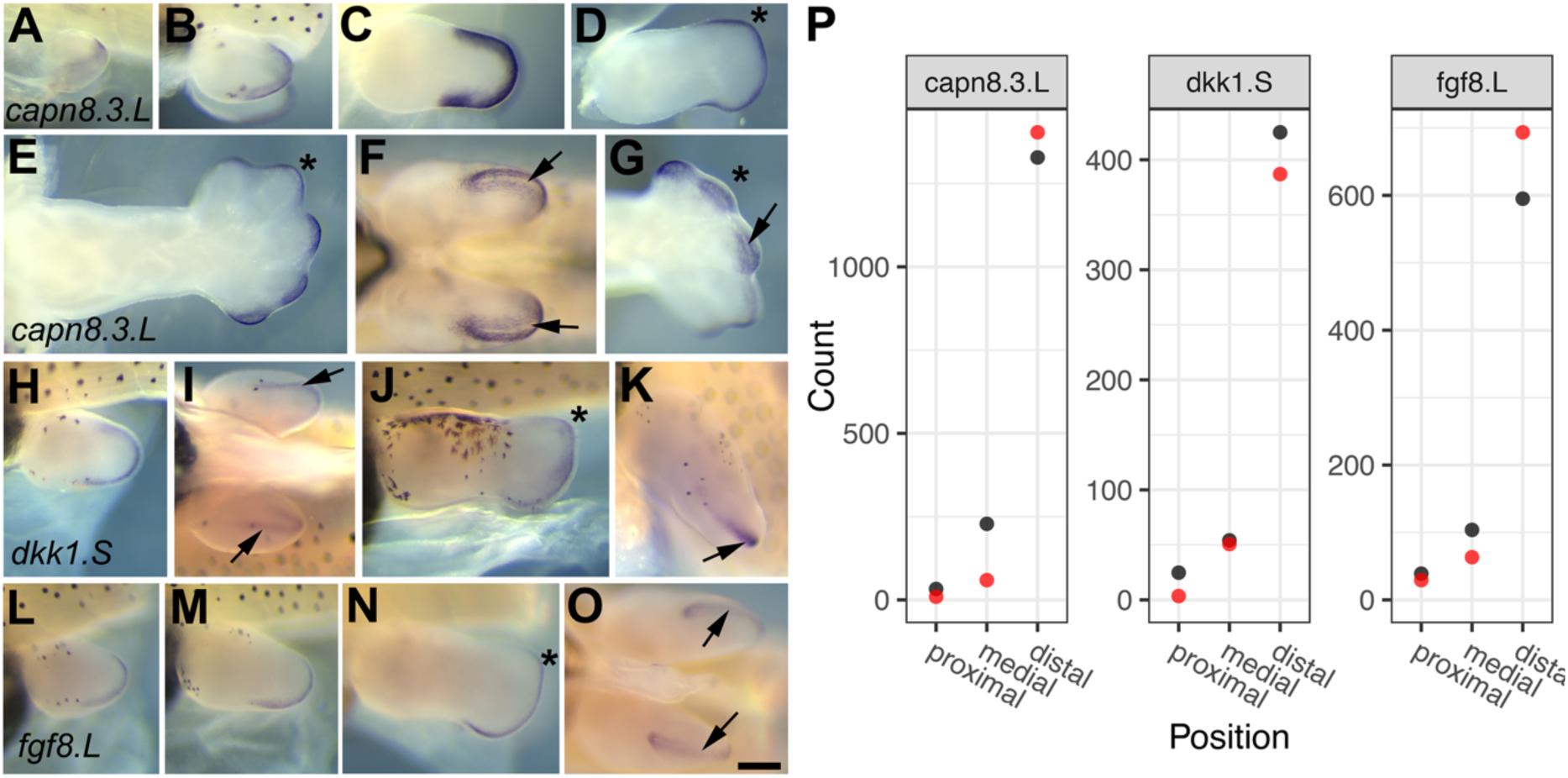
Expression of distal epithelial genes in *X. laevis* hindlimb buds. **A-O**, Whole mount in situ hybridisation, gene expression is marked by dark purple staining. **A-G** *Capn8*.*3L* is expressed in the epithelial cells flanking the AER. A-E Dorsal view of limb with distal to the right and posterior uppermost. **A**: stage 50, **B**: stage 51, **C**: stage 52, **D**: stage 53, **E**: stage 55 **F, G** views of the distal limb(s) to show the AER (black arrow), **F**: stage 53, **G**: stage 55. H-K expression of *dkk1*.*S* **H**: stage 51, dorsal view, **I**: distal view, **J**: stage 53 dorsal view, **K**: distal view. **L-O** expression of *fgf8*. **L**: stage 51 dorsal view, **M**, stage 52 dorsal view, **N**: stage 53 dorsal view, **O**: stage 53 distal view. Black arrows indicate the AER in distal views, and the approximate position of digit IV is indicated by an asterisk from stage 53. Black stellate cells are melanophores in non-albino tadpoles (B, H-M). Scale bar in O is 250 μM and applies to all panels. **P**, Normalised read counts of *capn8*.*3*.*L, dkk*.*S* and *fgf*.*S* in the three sections of the stage 51 limb bud transcriptome, red and black indicate biological replicate data.

Closer scrutiny showed that the AER margin cells do not express *capn8*.*3*.*L*, but rather it is expressed either side of the boundary, and appears stronger on the dorsal side. (Figure 3 F,G). *Capn8* is considered a classical calpain and *capn8* is expressed in the gastrointestinal tract of mammals (reviewed in Spinozzi et al 2021(Spinozzi et al., 2021)). In *Xenopus*, it was identified as a downstream target of Tfap2, expressed throughout the non-neural ectoderm in neurula stage embryos (Luo et al., 2005), so it may have a role in boundary definition. *Capn* genes have not been previously associated with vertebrate limb development. *Capn8* likely originated in the ancestor of jawed vertebrates (Macqueen and Wilcox, 2014) where it may have assumed a role in neural crest before being later co-opted into limb development. In *X. laevis, capn8* appears to have been duplicated many times: *capn8*.*1, 8*.*2, 8*.*3, 8*.*4 and 8*.*5* are clustered together with *capn2*.*L* on the 5L chromosome and *capn8*.*1 and 8*.*6* cluster with *capn2*.*S* on the 5S chromosome (data from *X. laevis* genome build v9.2 on Xenbase (Bowes et al., 2008)). In the diploid *X. tropicalis, capn8*.*1, 8*.*2, 8*.*3, 8*.*5 and 8*.*6* are clustered together with *capn2* on chromosome 5. *Capn8*.*2*.*L* was also a cluster 2 gene, mainly expressed in distal hindlimb samples, albeit at lower levels than *capn8*.*3L*, suggesting that they could be co-regulated.

Both homeologs of *dkk1* were cluster 2 genes. *Dkk1*.*S* (distal>proximal Log2 fold change 5.0), encodes dickkopf 1, an extracellular Wnt signalling antagonist that acts upstream of receptor activation. *Dkk1* was first discovered in *Xenopus* (Glinka et al., 1998) and is one of three dickkopf genes. In *Xenopus* hindlimbs, *dkk1* expression is AER-specific from stage 51 (Figure 3 H-K). Expression mimics that of *fgf8*, in the AER (Figure 3 L-O). This is very different from *dkk1* expression in the mouse limb bud, which is mesenchymal, and at the equivalent stage is expressed in two proximal spots, one posterior and slightly overlapping *shh* (ZPA) and one anterior (Grotewold et al., 1999, Monaghan et al., 1999), with later expression is in the interdigits. *Dkk1* knockout mice are non-viable and have fused and duplicated digits resulting from an expanded AER (Mukhopadhyay et al., 2001). In chickens, overexpression of *dkk1* leads to apoptosis of the AER cells and subsequent limb truncation (Grotewold and Ruther, 2002). While this clear difference in expression of *dkk1* in limb buds of amniotes (mesenchyme) and *Xenopus* (AER) is unexpected, a search for *dkk1* enhancers identified a conserved element in the 3’ UTR which was able to drive expression of a reporter very specifically in the developing mouse limb bud AER (Lieven et al., 2010). This element is conserved in fish, and therefore could also be present in *Xenopus*. Conversely, perhaps the mesodermal enhancer has been lost or masked in frogs.

The role of the fibroblast growth factor member Fgf8 in AER function and limb outgrowth in amniotes is well established (Lewandoski et al., 2000, Moon and Capecchi, 2000) and the expression of *fgf8* in the AER has also been well documented in *Xenopus* (Christen and Slack, 1997, Endo et al., 2000, Wang and Beck, 2014) Both *fgf8* homeologs were in cluster 2 and *Fgf8*.*L* (distal>proximal Log2 fold change 4.3) is shown here for comparison (Figure 3, L-O).

#### 2.3.2 Zic5.S and Sall1.S are expressed in the distal limb bud mesenchyme

Cluster 2 also contains several genes known to be preferentially expressed in the distal mesenchyme of amniote limb buds, including both homeologs of *fgf10*. Also in cluster 2 are 5’ hox transcription factors *hoxa13*.*S, hoxa13*.*L, hoxd13*.*S* and *hoxd13*.*L* (Figure 1E), previously shown to be distally expressed in *Xenopus* hindlimbs (Christen et al., 2003). Genes coding for several family members of the receptor tyrosine kinase/Fgf antagonist Sprouty (Spry) are also found in cluster 2 (*spry4*.*L, spry4*.*S, spry1*.*L, spry1*.*S* and *spry2*.*L)*. We have previously shown that *X. laevis spry2* was expressed in limb bud AERs, and *spry1* and *spry 4* in limb bud mesenchyme (Wang and Beck, 2014).

Two genes encoding transcription factors found in with cluster 2 that were highly significantly enriched in distal stage 51 limb bud (Figure 1E) were validated and further analysed using whole mount *in situ* hybridisation of *Xenopus* hindlimbs. Mutations in the gene coding for the spalt-like zinc finger transcription factor Sall1 are associated with Townes-Brocks syndrome (OMIM #107480). The phenotype includes extra thumbs, radial dysplasia and foot deformities (Townes and Brocks, 1972) which suggests a role in anterior-posterior limb axis patterning of the autopod and zeugopod. *Sall1* is expressed in stage E10.5 mouse limb buds and has a distal mesenchyme and AER expression pattern in early chicken limb buds (Buck et al., 2001, Farrell and Munsterberg, 2000). *Sall1*.*S* reads were almost exclusively found in the distal hindlimb samples (distal>proximal log2 fold change 5.3). Expression of *sall1*.*S* was localised to the distal mesenchyme at stage 50 and 51 (Figure 4 A,B,E), but from stage 52 the expression was lost from the most distal cells, and was stronger anteriorly and dorsally (Figure 4 C, D, F). *Xenopus* has four *spalt-like* orthologues: *sall1*.*L sall3*.*L* and *sall3*.*S* were also cluster 2 genes, *sall4*.*L and sall4*.*S* were found in cluster 1. Previous work by Neff et al revealed the role of *sall4* in *Xenopus* limb regeneration, and stage 53 control limbs appear to have similar expression to *sall1* here (Neff et al., 2011). *Sall2* homeologs were not represented in any of the clusters, and read numbers were very low suggesting *Sall2* is not a key player in limb developmental patterning in early *Xenopus* limbs

**Figure 4:**
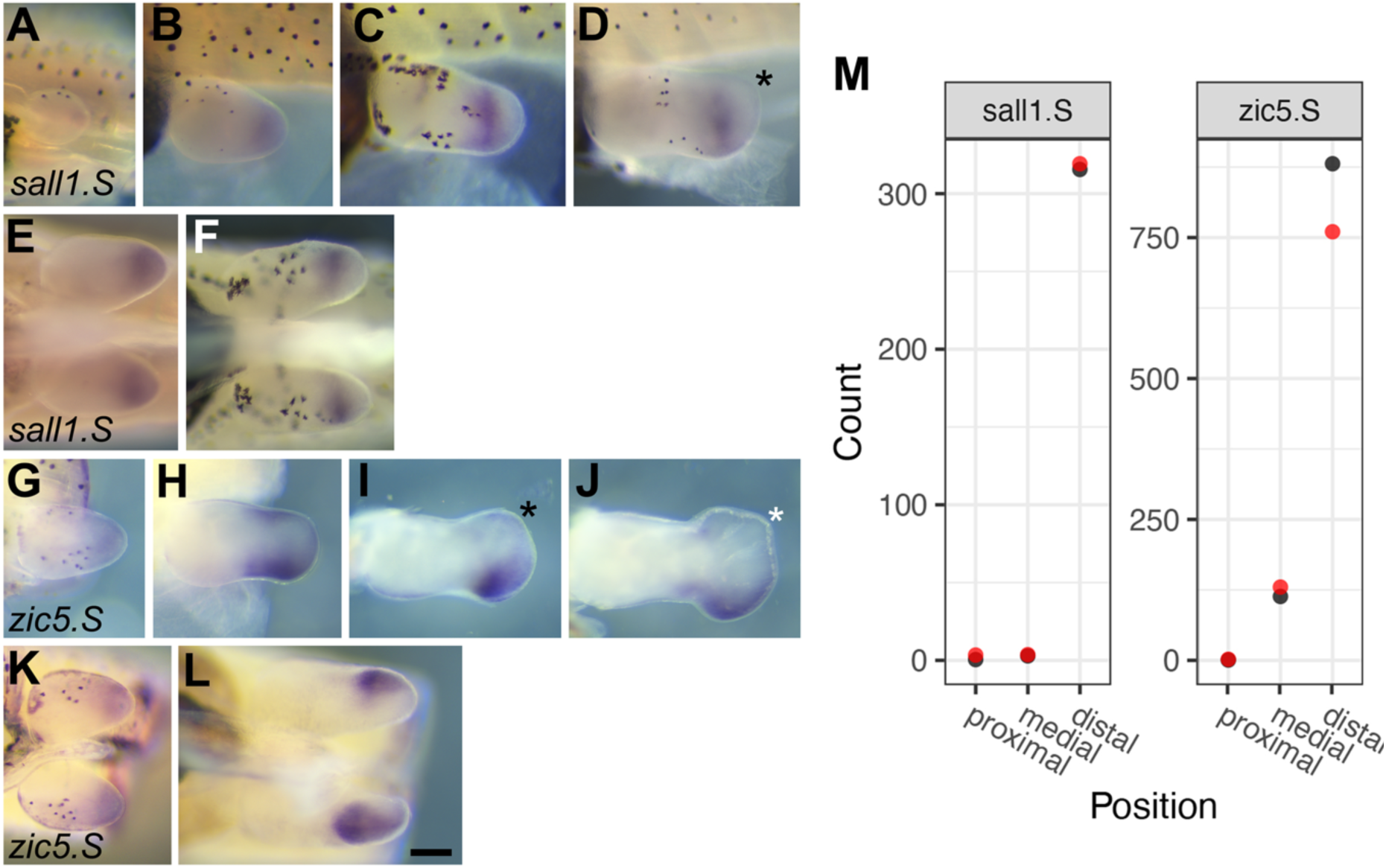
Expression of two distal mesenchymal genes in *X. laevis* hindlimb buds. **A-L** Whole mount in situ hybridisation where expression is indicated by dark purple staining. **A-F** expression of *sall1*.*S*. **A-D** dorsal views of the limb with distal to the right and posterior uppermost. **A**: stage 50, **B**: stage 51, **C**: stage 52, **D**: stage 53. **E-F** views of the distal limb, the ventral midline of the tadpole running left to right in the centre of the image. **G-L** expression of *zic5*.*S* indicated by dark purple staining. **G-J** dorsal views of the limb. **G**: stage 51, **H**: stage 52, **I**: stage 53, **J**: stage 54. **K** and **L** views of the distal limb, **K**: stage 51, **L** stage 54. Asterisk indicates the approximate location of digit IV in stage 53 and above limb buds. Melanophores are present on some limb buds (A-D, F, G, K) as black stellate cells. Scale bar in L is 250 μM and applies to all panels. **M**, Normalised read counts of *sall1*.*S and zic5*.*S* in the three sections of the stage 51 limb bud transcriptome, red and black indicate biological replicate data, where only red can be seen it overlaps completely with black.

The zic family of zinc finger transcription factors are orthologues of *Drosophila odd-paired*, with tetrapods, including *Xenopus*, normally have five family members (Aruga et al., 2006, Houtmeyers et al., 2013). Cluster 2 contains several distally located *zic* genes (*zic1*.*S, zic2*.*L, zic2*.*S, zic3*.*L, zic3*.*S, zic5*.*L* and *zic5*.*S*). *Zic1* and *zic4* form a tandem pair on chromosomes 5S and 5L, homeologs have very few, mostly distal reads in limb buds. *Zic3* is a singleton in tetrapods, and is expressed distally, and *zic2* and *zic5* form a tandem pair on chromosomes 2S and 2L, and have much higher read counts. *Zic2* has been previously shown to be expressed in distal limb bud mesenchyme in mice(Pan et al., 2011), but *Zic5* has no known association with limb development. *Zic5*.*S* (distal>proximal log2 fold change 3.9) expression was validated using whole mount limb in situ hybridisation (Fig 4 G-L). Expression was seen very faintly in distal mesenchyme at stage 51, (Figure 4 G, K) and in two regions at stage 52, a broad expression in the distal anterior and a more restricted stripe in posterior mesoderm (Figure 4H). After stage 52, the expression in posterior mesenchyme was lost, with anterior/dorsal expression maintained in the forming autopod at stage 53-54 (Figure 4 I, J, L).

### 2.4 Genes with differential expression across the proximodistal axis were associated with cell adhesion, cell proliferation, as well as Wnt, Fgf and retinoic acid signalling

We annotated 894 genes with differential expression of log fold change +/->1.5 between proximal and distal limb bud samples, and analysed these for overrepresented gene ontologies at all levels, compared to all *X. laevis* annotated genes (9414). The complete data for 36 overrepresented GO terms, at all levels, can be found in the supplementary data file, but we will highlight the most pertinent ones here.

#### 2.4.1 Cell adhesion

Differential cell adhesion has been shown to correlate with proximodistal position in early limb buds (Wada, 2011). In the *Xenopus* stage 51 hindlimb, 56 genes with differential proximodistal expression were associated with the GO term “cell adhesion”. Nineteen of these had significantly more reads in the distal third and 34 had more in the proximal third (three genes were removed due to having low read numbers) (Figure 5). Of particular interest is *Agr2*.*L*, which codes for a thioredoxin domain, secreted, GPI anchored cell surface protein. Agr2 was originally identified as nAG (newt anterior gradient), which is a ligand for the salamander Prod1 receptor, which plays a role in establishing proximodistal identity in limb regeneration (da Silva et al., 2002).

**Figure 5.**
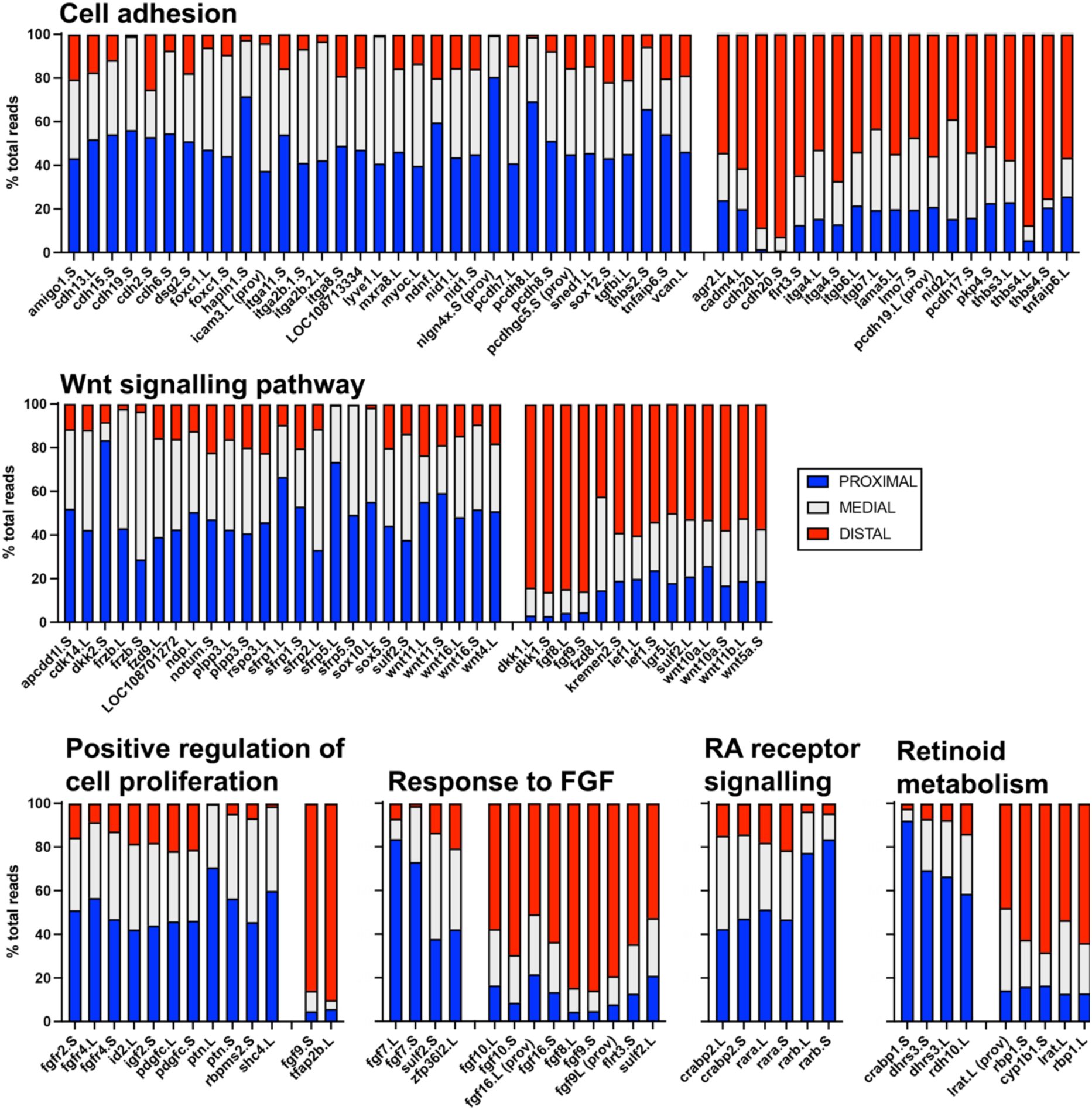
Relative distribution of read counts for genes in differentially expressed gene ontology gene sets for cell adhesion, Wnt signalling pathway, positive regulation of cell population proliferation, response to fibroblast growth factor, retinoic acid receptor signalling and retinoid metabolism. Stacked graphs show the percentage of proximal reads is in blue, medial reads in grey and distal in red. Genes with statistically more reads in distal than proximal are on the left and those more proximal on the right, and are ordered alphabetically within each group.

Additionally, we identified proximodistal differential expression of genes encoding members of the integrin, cadherin, thrombospondin and nidogen families. Six genes encoding integrins, which mediate cell to matrix adhesion, were differentially expressed, with those coding for subunits α8, α11, α2b found at highest levels in the proximal limb bud and α4, β6 and β7 found at highest levels distally. Cadherins (Cdh) are calcium-dependent transmembrane molecules involved in cell to cell adhesion. *Cdh20* was almost exclusively expressed distally, whereas *cdh2* (N-cadherin), *cdh6, cdh13, cdh15* and *19* were highest in the proximal segment. The cadherin-related desmoglein *dsg2* is found proximally. Protocadherins *pcdh19* and *17* are distal, whereas *pcdhgc5* is proximal.

Thrombospondins (*thbs1* to *4*) are calcium binding, extracellular matrix proteins implicated in skin wound healing in mammals. In *X. laevis* limb buds, *thbs*3 and 4 are distal (cluster 2) and *thbs2* is proximal (cluster 6S/4L). Thrombospondins have been previously implicated in axolotl limb development and regeneration (Whited et al., 2011) and *X. laevis* limb regeneration (Pearl et al., 2008). The basement membrane protein Nidogen (Ndg) has two members, *ndg1* is proximal, whereas *ndg2* is distal. In mice, the two nidogens appear redundant, but double knockouts have soft tissue syndactyly resulting from failed interdigital apoptosis (Bose et al., 2006).

#### 2.4.2 Wnt agonists and antagonists are both proximally and distally expressed

The Wnt signalling pathway is most often associated with dorsal-ventral limb patterning, especially the ligand Wnt7a, which helps establish the position of the AER at the dorsal ventral boundary and restricts *lmx1b* to dorsal mesenchyme (Riddle et al., 1995). Here, we found Wnt pathway genes were significantly over-represented in our proximal-distal analysis. *Dkk1*.*L and* .*S*, which code for Wnt pathway antagonists, were the most distally biased genes in this set, are captured in cluster 2, and the AER-specific expression of *dkk1*.*S* has been previously discussed (Figure 3). Interestingly, *Dkk2*.*S* shows the opposite polarity, with over 80% of transcripts on the proximal limb bud. In *Xenopus* embryos, Dkk2 acts as an activator of canonical Wnt signalling rather than an antagonist (Wu et al., 2000): so both expression and function of these two *dkk* genes are in opposition. *Dkk2*.*S* is a cluster 4 gene, but its homeologue *dkk2*.*L* falls into cluster 7, as does another family member, *dkk*.*3*.*S*. Cluster 7 genes have highest expression in the medial limb bud, which would indicate that there is a Dkk associated with each of the three proximodistal domains in *Xenopus* limbs. Interestingly in mice, *dkk1*, 2 and 3 are all mesenchymal in developing limb buds, with complementary expression patterns around the time of mesodermal condensation formation(Monaghan et al., 1999). *Xenopus* Dkk3 does not appear to modulate Wnt signalling (Wu et al., 2000) and instead affects mesoderm induction via interference with TGFβ pathways (Pinho and Niehrs, 2007).

Genes encoding the ligands Wnt4, 16 and 11 are more proximal and Wnt10a, 11b and 5a are more distal. Wnt5a is associated with some cases of autosomal dominant Robinow syndrome, a rare form of dwarfism with shortening of the mid and distal parts of the limbs. It is expressed in the mesoderm just under the AER, often referred to as the progress zone, in chicken limb buds (Dealy et al., 1993). The same expression is seen in axolotl and mouse limb buds, and *wnt5a* null mice have distally truncated limbs (Parr et al., 1993, Lovely et al., 2022). The effect of Wnt5a on limb proximodistal patterning is mediated through the planar cell polarity pathway rather than canonical Wnt signalling and is dependent on AER Fgf signals (Gao et al., 2018). Notum is a Wnt pathway antagonist, with expression highest in the distal limb bud. Expression in the early chicken limb was previously shown to be mainly in the AER (Saad et al., 2017). In contrast, secreted frizzled related protein genes, encoding antagonists known as Sfrps, are predominantly expressed in the medial and proximal limb bud. Sfrp are able to antagonise Wnt by mimicking receptors and binding Wnt, but may also have other non-Wnt roles (for review see Nathan and Tzahor, 2009).

#### 2.4.3 Cell proliferation and response to fibroblast growth factor (Fgf)

FGF pathway agonists and antagonists are both proximally and distally expressed in stage 51 *Xenopus* hind limb. The role of AER-Fgf signalling in proximodistal vertebrate limb bud patterning is well established, with Fgf8 as the key functional molecule (Lewandoski et al., 2000, Moon and Capecchi, 2000). We have already discussed *fgf8* (see Fig 3). Amniotes have additional Fgfs in the AER, such as Fgf4, 9 and 17 (Martin, 1998). *Fgf4* transcripts were not detected in *Xenopus* hindlimb buds at stage 51, however, *fgf9, fgf10* and *fgf16* are additional distal Fgfs (all in cluster 2). Conversely, *fgf7* is proximal, S/L forms are found in cluster 6/4 respectively.

Most of the genes associated with the GO term cell proliferation are located predominantly in the distal limb, including transcripts for growth factors Igf2 and Pdgfc and Fgf receptor coding genes *fgfr2* and *fgfr4*. The remaining two Fgf receptor genes are expressed at similar levels in all three proximodistal regions of the stage 51 limb bud (see supplementary data file). Over 90% of transcripts for cluster 2 genes *tfap2b* and *fgf9* are in the distal limb. Oddly, the two *X. laevis* homeologs of *sulf2*, encoding heparan sulphate 6-O-endosulfatase enzymes, are expressed in different regions, with the S-form proximal and the L-form distal, as previously confirmed by in situ hybridisation (Wang and Beck, 2015). Sulf2 was also captured in the Wnt signalling GO.

#### 2.4.4 Retinoic acid metabolism and signalling

Retinoic acid (RA) is proposed as a proximal signal opposing distal Fgf to establish expression domains across forming limb buds (Mercader et al., 2000). Two GO terms associated with RA were found to be significantly associated with proximal vs. distal gene expression in our stage 51 hindlimbs (Figure 5). Retinoic acid receptor signalling genes were all proximally biased, and included the genes for retinoic acid receptor subunits A and B (*rara, rarb*), and the binding protein Crabp2 (previously characterised in McEwan et al., 2011). *Rarb* showed particularly strong proximal bias, and a third RA receptor-encoding gene, *Rarg* was not differentially expressed (see supplementary data file). Genes associated with retinoid metabolism were also over-represented, with both proximal and distally biased expression (Figure 5).

### 2.6 Retinoic acid down-regulated genes are expressed in the distal hindlimb segment

Retinoic acid (RA) has been demonstrated to reprogram distal cells to a more proximal fate in chicken limb bud grafts (Tamura et al., 1997), and exposure to RA in regenerating frog limb buds (Maden, 1982) and axolotl limbs (Monaghan and Maden, 2012) results in proximodistal duplications, suggesting RA disrupts pattern along this axis. RA is produced endogenously from vitamin A and can modulate gene transcription via binding to RA nuclear receptors that recognise retinoic acid response elements (RARES) in enhancers (Cunningham and Duester, 2015). RA reporter activity has been detected in forelimbs, but not hindlimbs, during axolotl development (Monaghan and Maden, 2012). We previously showed the presence of opposing RA synthesis and catabolic genes in early developing *Xenopus* limb buds. To see if there was any evidence from RA target genes to support a functional role for RA gradients in the early limb bud, we looked at the proximodistal distribution in *Xenopus* stage 51 hindlimb of genes identified as up- or down-regulated by RA treatment in a prior study of axolotl limb regeneration (Nguyen et al., 2017). Nyugen et al identified 101 genes as up-regulated by RA, 14% of these had no recognised homologues. Of the remaining genes, 62% were not significantly differentially expressed across *Xenopus* stage 51 limb buds, and a further 7% were not expressed in limb buds at all. Twenty-seven genes (31%) were differentially expressed (Figure 6) with all clusters represented. Proximally expressed genes were most common (18 genes, clusters 4, 5 and 6) followed by distal genes (13 genes, clusters 1 and 2) and 5 genes were predominantly expressed in the medial region (cluster 7) with one gene lowest in the medial region (cluster 1). Therefore, although many salamander RA up-regulated genes had differential expression across the proximodistal axis in stage 51 frog hindlimb buds, there was no consistent pattern observed. However, *meis* genes were consistently found in cluster 4 (proximal genes), as expected from previous work (Mercader et al., 2000), and the related homeobox gene Pbx1 was in cluster 6 (proximal to distal gradient). Our findings of proximal-biased expression for these genes therefore support the role of Meis homeobox transcription factors as RA-regulated mediators of proximal identity. The finding that conditional inactivation of *meis1/2* in mice results limb phocomelia, where proximal elements were missing but distal ones develop normally, further demonstrates the importance of RA regulated genes in limb proximodistal patterning (Delgado et al., 2020).

**Figure 6:**
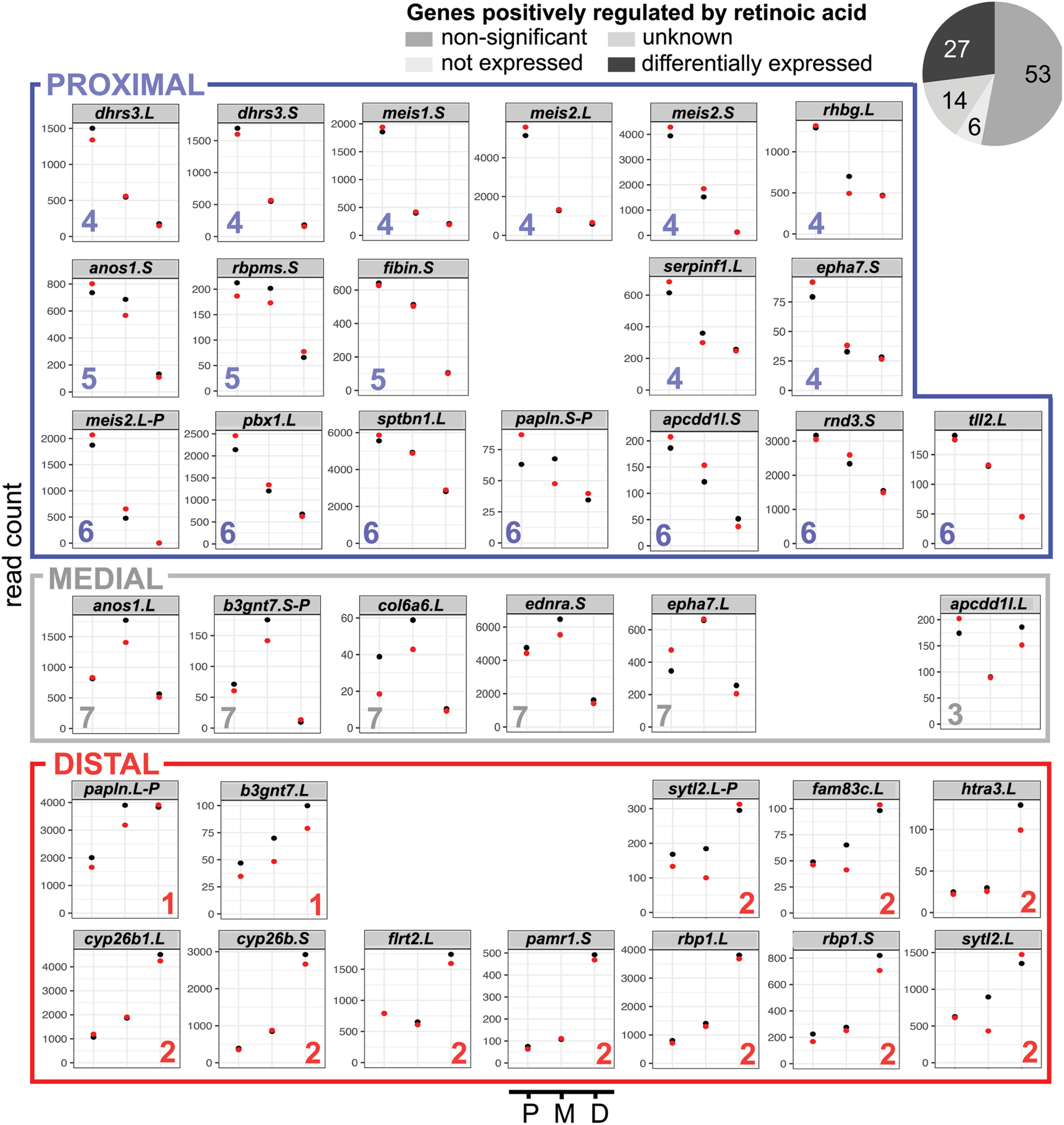
Genes previously shown to be up regulated by retinoic acid in axolotl limbs do not show consistent expression distributions in *Xenopus* limbs. 100 different genes identified as up-regulated by RA in axolotl limb (Nguyen et al., 2017) were identified one of four categories: unknown (no *Xenopus* homologue), not expressed (not found in *Xenopus* stage 51 limb bud), non-significant (expression is not significantly different across the three limb bud sections) or differentially expressed. Homeologs of the 27 differentially expressed genes in the allotetraploid *Xenopus laevis* are found in all seven clusters, cluster number is indicated by coloured number at the base of each graph, read counts are shown on the y-axis and position on the x-axis. P= proximal, M=medial, D=distal. Replicates are represented by red and black dots and where only one dot can be seen, these overlap completely. Coloured boxes are used to group clusters with proximal, distal or medial/terminal expression patterns.

As well as promoting the proximal to distal spatial pattern via upregulation of Meis homeobox proteins, it is possible that RA might also play a role in limiting the expression of distally localised genes via inhibition. Interestingly, of the 14 genes identified in axolotl limb regeneration as down-regulated by RA(Nguyen et al., 2017), ten (72%) were differentially expressed across the early *Xenopus* limb buds with all of these falling into cluster 2 indicating predominantly distal expression (Figure 7). A further three genes were present in limbs but not differentially expressed, and one gene was not expressed in limb buds. While distal expression is indicated by inclusion in cluster 2, *hoxa13, lhx2, rgs18, rgs2, spry1 and zic5* had expression that was predominantly distal, whereas *dner* and *lhx9* had more graded distal to proximal expression and *lmo1* and *msx2*, which both encode transcription factors, were least expressed in the medial region (Figure 7).

**Figure 7:**
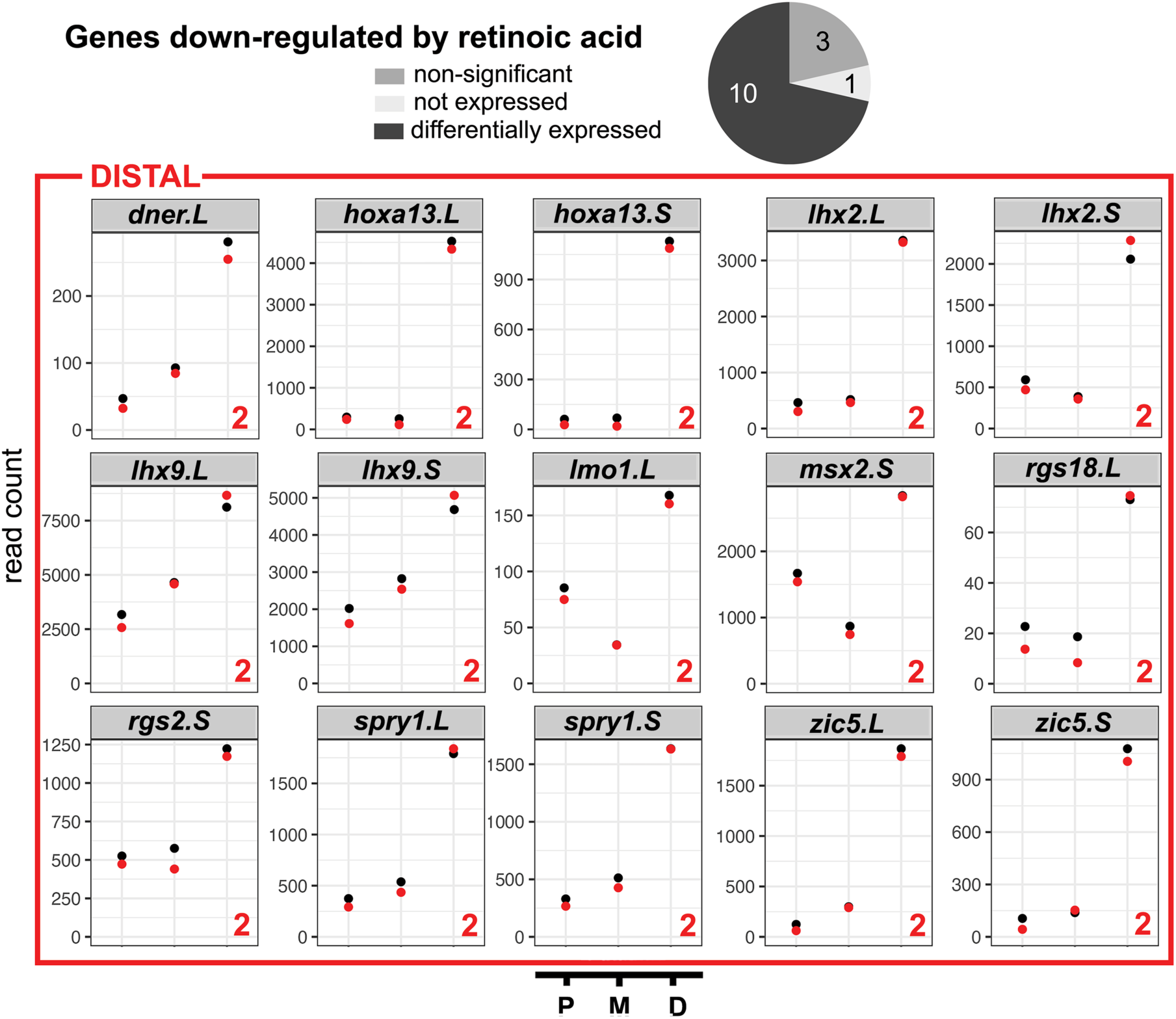
Distal gene expression in stage 51 *Xenopus* hindlimb bud of genes previously shown to be down regulated by retinoic acid in axolotls. Pie chart: 14 different genes identified as down regulated by RA in axolotls (Nguyen et al., 2017) were identified as not expressed (not found in *Xenopus* stage 51 limb bud), non-significant (expression is not significantly different across the three limb bud sections) or differentially expressed. Homeologs of the 10 differentially expressed genes in the allotetraploid *Xenopus laevis* are found exclusively in cluster 2. Cluster number is indicated by coloured number at the base of each graph, read counts are shown on the y-axis and position on the x-axis. Replicates are represented by red and black dots and where only one dot can be seen, these overlap completely. P= proximal, M=medial, D=distal. Coloured box indicates all genes had distal expression patterns.

## 3. Conclusions

Our analysis of gene expression across the proximal to distal axis of a stage 51 tadpole hindlimb bud provides a an unbiased overview of the genes and mechanisms that underly the patterning of the *Xenopus* limb bud at this critical stage of its development. Stage 51 was chosen based on our previous work proposing a transient, limb bud-autonomous, proximodistal RA gradient(McEwan et al., 2011). We show that the distal segment, which incorporates the cryptic AER, is the most transcriptionally distinct, and describe the novel expression of a calpain encoding gene, *capn8*.*3*.*L* in the flanking regions of the AER. We also identify and describe expression of three further genes previously unlinked to limb development, tightly focused in either the proximal (*tnn*.*L, gata5*.*L* or distal mesenchymal (*zic5*) region of developing *Xenopus l*imbs. Gene ontology analysis of genes expressed differentially between proximal and distal segments provided support for differential cell adhesion, proliferation, Wnt and Fgf signalling as well as RA signalling, although confirmation of these results is beyond the scope of this study. There was no clear pattern to Wnt or Fgf pathway components, with agonists and antagonists for both pathways found both proximally and distally, suggesting regulation of these pathways is complex and nuanced. In contrast, we saw clear evidence for retinoic acid signalling. All genes associated with RA receptor signalling were proximal, and all the genes identified as down-regulated by RA in a previous study of axolotl limb regeneration(Nguyen et al., 2017) were distally expressed in *Xenopus* early hindlimbs. This suggests RA signalling is limited to proximal cells and that RA is actively cleared from distal limb buds. Our findings show support for the model of positional identity conferred by proximal RA and distal Fgf signals in early limb buds proposed by others(Mercader et al., 2000, Saiz-Lopez et al., 2015).

## 4. Experimental Procedures

### 4.1 Animal procedures

All animal procedures were approved by University of Otago’s Animal Ethics Committee as AEC56/12. *Xenopus* laevis adult females were injected with 500U HCG per 75g bodyweight, into the dorsal lymph sac, to induce egg laying. Eggs were fertilised *in vitro* using fresh testis preparations and jelly removed with 2% cysteine HCl. From stage 47, tadpoles were fed a suspension of spirulina powder and housed in tanks in a recirculating aquarium.

### 4.2 Sample preparation

Stage 51 hindlimb buds were dissected into distal, medial and proximal thirds, each with a length of approximately 200 μM, using Vannas iridectomy scissors. To do this, six tadpoles at a time were anaesthetised with 1/4000 v/v MS222 (tricaine) in 0.1 x MMR (Marc’s modified Ringers: 10 mM NaCl, 0.2 mM KCl, 0.1 mM MgSO_4_.6H_2_O, 0.2 mM CaCl_2_, 0.5 mM HEPES, 10 μM EDTA, pH 7.8). The tadpoles were placed onto damp Whatman paper and the distal third of the hindlimb cut off and placed in 1.5 ml sterile Eppendorf tube cap containing 50 μl of lysis buffer, using sharpened Dumont #5 forceps. When all six distal samples had been collected the cap was placed on a tube containing lysis buffer, spun briefly to pellet the tissue, and placed on dry ice. The process was then repeated for medial sections, and finally for proximal sections by detaching the remaining limb tissue from the body wall cleanly. The tadpoles were then rotated onto the other side and the process repeated with fresh caps. The samples for each region were combined by spinning the new caps with the original tubes. Two different sibships of tadpoles were used, and separately prepared “red” and “black” biological replicates from these were each comprised of 30 pooled limb slices per sample.

Distal, medial and proximal samples for each replicate were homogenised with an OMNI TH handheld 5 mm tip probe in 400 μl lysis buffer (Qiagen) for 20 seconds. RNA was extracted using a Qiagen RNeasy mini kit and on-column DNAse I according to the manufacturer’s instructions.

### 4.3 Transcriptome sequencing, processing, read mapping and quantification

Illumina TruSeq RNA Sample Prep Kit was used with 1 μg of each total RNA for the construction of sequencing libraries using standard Illumina protocols. Paired-end sequencing reads were generated by Illumina HiSeq at the Otago Genomics Facility, and data provided as FastQ files. Raw data has been deposited in NCBI Gene expression omnibus (GEO) as GSE179158. Illumina adapter sequences were trimmed from reads and low-quality quality sequences were removed using trimmomatic v0.39 (Bolger et al., 2014). HISAT2 v2.2.1 (Kim et al., 2015) was used to align reads for each sample. A HISAT2 index was built from the *X. laevis* transcriptome, genome, and gene models from Xenbase v9.2 (Karimi et al., 2018) using the “hisat2-build” command. Samples were aligned using the “hisat2” command, converted to BAM files and sorted using samtools v1.10) (Li et al., 2009). Sorted files were quantified using the featureCounts v 2.0.1 (Liao et al., 2014) which returned a combined quantification file for all samples.

### 4.4 Differential expression

The combined quantification file from featureCounts was loaded into R using read.table. The table was made into a DESeq object using the DESeq2 (Love et al., 2014) command “DESeqDataSetFromMatrix” (options: colData = data.frame(position = factor(position)), design = ∼ position), where position identified what portion of the limb bud a sample came from. The resulting DESeq object was filtered for transcripts with low counts (counts >= 10). The counts table output was made to be a DESeq object with DESeq2’s command “DESeqDataSetFromTximport” (options: colData = samples, design = ∼ position). This DESeq object was also filtered for transcripts with low counts (counts >= 10). Differential expression analysis was performed with the DESeq2 package. Conditions (proximal, medial and distal segments of limb) were compared using DESeq2’s Wald test. The command “DESeq” was used for this with default options. Differential analysis results were filtered for an adjusted p-value < 0.05, and log2 fold change +/-1. Full data can be found in the supplementary data file.

### 4.4 Hierarchical read clustering and volcano plot

For each limb segment, the normalised counts from replicates were averaged, and the resulting matrix clustered using the package “ComplexHeatmap” (Gu et al., 2016) using the command “Heatmap, option: row_km = 7”. Rows were clustered using Pearson distance. A full list of genes in each cluster can be found in the supplementary data file. Volcano plots were made using VolcaNoseR (Goedhart and Luijsterburg, 2020).

### 4.5 In situ hybridisation

*Gata5*.*L, irx5*.*L, tnn*.*L, sall1*.*S, zic5*.*S, capn8*.*3*.*L* and *dkk1*.*S* were amplified from reverse transcribed limb bud cDNA and blunt cloned into the EcoRV site of pBSIIKS+. Primers can be found in Table 1. *Fgf8* probe was previously described (Pownall et al., 1996) with modifications for limb buds from(McEwan et al., 2011). Images were captured using a Leica FluoIII microscope using Leica LS software, with embryos submerged in PBSA on a 2% noble agar lined petri dish and LED lighting. Adobe photoshop CC 2019 was used to assemble figures.

**Table 1:**
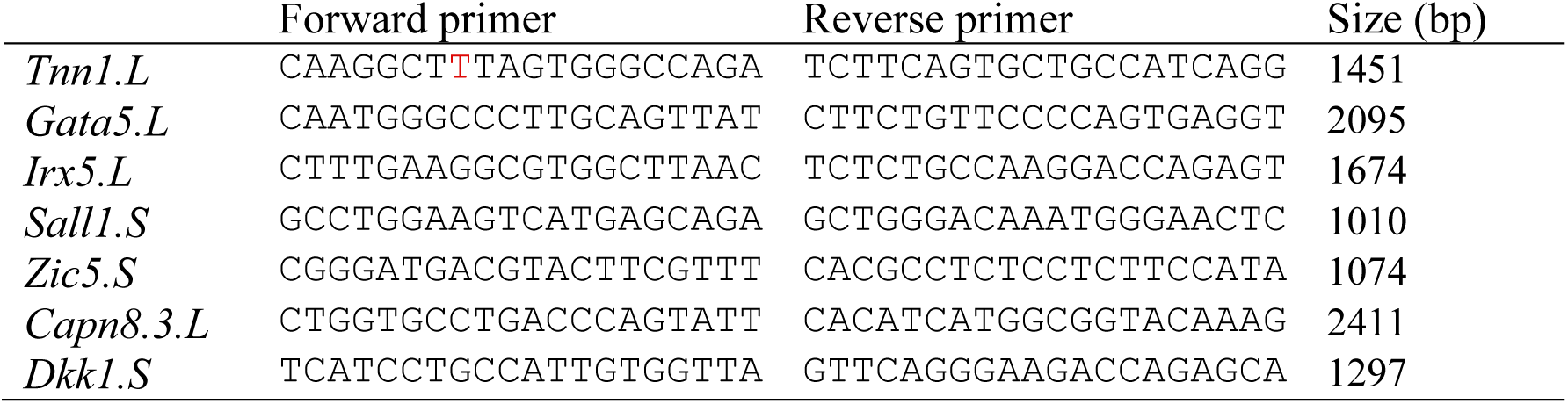
Primers used to amplify cDNA for *in vitro* transcription, to generate in situ hybridisation probes.

### 4.6 GO Analysis

Gene ontology (GO) analysis was performed on significantly differently expressed genes for the proximal vs. distal contrast, with a cut-off of log2 +/-1.5. The clusterProfiler package (Yu et al., 2012) and the *X. laevis* genome wide annotation database (Carlson, 2019) were used for this. Prior to analysis, gene names were annotated with Entrez IDs using the “GenePageLaevisEntrezGeneUnigeneMapping.txt” file from Xenbase. The “enrichGO” command (options: OrgDb = org.Xl.eg.db, ont = “BP”, readable = TRUE) provided enrichment analysis for the genes contained in the proximal vs. distal set. All GO levels were included, and 36 terms were significantly over-represented (PAdj <0.05). Full data for GO can be found the supplementary data file.

### 4.7 Analysis of Retinoic acid regulated genes

Supplemental data from (Nguyen et al., 2017) that identified genes up and down regulated in axolotl limb regeneration supplemental table was manually cross-referenced with DE genes in this study. Genes that did not map to one of the seven clusters, or that had fewer than 100 reads across all samples were eliminated. Read counts of genes both differentially expressed across the proximodistal axis of the *Xenopus* limb bud and regulated by RA in axolotls were plotted for proximal, medial and distal using ggplot2 (Wickham, 2016) in R and were grouped according to cluster.

### 4.8 Data availability

Raw RNA-Seq fastq files, the HISAT2/featureCounts table, and a file of significant differential expression is available at NCBI GEO with accession GSE179158. Read counts, differential expression, a list of genes in each cluster, and gene ontology terms can be found in the supplementary data file.

## Acknowledgements

This work was supported by a University of Otago Research grant to CWB and by the Department of Zoology. We are grateful for the excellent support and maintenance of our *Xenopus* colony provided by Nikita Woodhead. Sequencing was carried out at the Otago Genomics Facility.

DTH: formal analysis, data curation, visualisation. JSB and JMW: molecular cloning, in situ hybridisation. RCD: library preparation and sequencing, editing. TM: formal analysis. CWB: conceptualisation, sample collection and RNA extraction, writing, editing, visualisation and supervision.

